# Systematic beach monitoring as a health assessment tool: cetacean morbillivirus under non-epizootic circumstances in stranded dolphins

**DOI:** 10.1101/2021.02.13.431109

**Authors:** Victor H. B. Marutani, Flávia Miyabe, Alice F. Alfieri, Camila Domit, Andressa M. R. N. de Matos, Mário R. C. M. Filho, Ana P. F. R. L. Bracarense

## Abstract

Cetacean morbillivirus (CeMV) was identified as the etiologic agent of several epizootic episodes worldwide. Most of these studies are based on unusual mortality events or identification of new viral strains. We investigated the occurrence of CeMV under non-epizootic circumstances at a world heritage in Southern Brazil by a combination of pathologic, immunohistochemical and molecular assays. From 325 stranded cetaceans, 40 were included. Guiana dolphin (*Sotalia guianensis*) was the species most frequent. Interstitial pneumonia and non-suppurative encephalitis were the main pathologic findings associated to CeMV infection. Intracytoplasmic immunolabeling anti-CeMV was observed mainly in lungs and lymph nodes. All samples were negative in RT-PCR assay. Diagnosis of CeMV is challenging in areas where epizootic episodes have not been recorded and due to *postmortem* changes. We observed a CeMV prevalence of 27.5%. The results described here increase the knowledge about CeMV under non-epizootic conditions in Brazil.

**Article Summary Line:** We observed a prevalence of 27.5% of CeMV in a World Heritage site of Paraná’s coast. The results indicate an increase in the prevalence of CeMV at this region and, possibly, a degradation of marine ecosystem. Marine mammals are sentinels of marine environment and the ocean health is inextricably linked to human health on a global scale.

## Introduction

Cetacean morbillivirus (CeMV) was first reported in cetaceans in an epizootic episode affecting more than 100 striped dolphins (*Stenella coeruleoalba*) in the Catalonian coast of Spain due to a combination of histopathologic findings, immunoperoxidase assay and virus isolation (*1, 2*). Since then, several unusual mortality episodes affecting different species of cetaceans have been linked to CeMV infection in Atlantic coast of United States of America (*3, 4*); Mediterranean Sea (*5–7*), and recently in southeastern Atlantic coast of Brazil where more than 250 Guiana dolphins (*Sotalia guianensis*) died (*8*).

CeMV species comprises three well characterized viral strains: the porpoise morbillivirus (PMV) (*9*), the dolphin morbillivirus (DMV) (*1, 10*), and the pilot whale morbillivirus (PWMV) (*11*). Three new CeMV strains were detected by reverse transcription polymerase chain reaction (RT-PCR) in Longman’s beaked whale (*Indopacetus pacificus*) from Hawaii (*12*), in IndoPacific bottlenose dolphins (*Tursiops aduncus*) from Western Australia (*13*), and in Guiana dolphin (*Sotalia guianensis*) from Brazil (*14*). The latter strain was linked to an unusual mortality event in Rio de Janeiro, Brazil, due to a combination of pathologic and molecular evidence and named Guiana dolphin-CeMV (GD-CeMV) (*8*). Recently, a novel strain similar to GD-CeMV was detected by RT-PCR in Southern right whales (*Eubalena australis*) from Brazil (*15*).

The detection of novel viral strains and their potential to cause unusual mortality events increases the concern about the preservation and conservation of cetaceans worldwide. Available data concerning cetacean morbilliviruses are mainly related to epizootic episodes or identification of new strains in a few individuals. Few studies regarding non-epizootic (NE) circumstances of CeMV worldwide were found when public databases were consulted, and none was accomplished in Brazil’s coast. Thus, the aim of this study was to investigate the occurrence of CeMV and other pathogens at a world heritage in Southern Brazil in NE circumstances and to describe the epidemiological, histopathological, immunohistochemical, and molecular findings.

## Study area

The study was accomplished in the coastline of Paraná state, Southern Brazil, comprising the Paranaguá estuarine complex (PEC) and the adjacent 100 km coastline (25°44’S and 48°29’W) within the PEC includes five major bays – Antonina, Guaraqueçaba, Laranjeiras, and Pinheiros – and three islands – Mel, Superagui, and Peças – encompassing a total area of 618 km^2^. The area is classified as a biosphere reserve and a World Heritage site by the United Educational, Scientific, and Cultural Organization (UNESCO).

The ocean beaches and PEC were monitored by the Projeto de Monitoramento de Praia da Bacia de Santos (PMP-BS), a requirement set by the Brazilian Institute of the Environment (IBAMA) for the environmental licensing of the oil and natural gas production and transport by Petrobras at the Santos Basin pre-salt province (25°05’S 42°35’W to 25°55’S 43°34’W) – for any stranded cetaceans. A field permit was granted by the Ministry of Environment-MMA (SISBIO 640/2015).

The research institutions participating in the study were the Laboratory of Animal Pathology-LAP and Laboratório de Virologia Animal-LabViral (Universidade Estadual de Londrina-UEL), and Laboratório de Ecologia e Conservação-LEC (Universidade Federal do Paraná/Centro de Estudos do Mar-UFPR).

### Animals

All carcasses were referred to LEC-UFPR where data such as gender, development state, measurement of total length, weight, and *postmortem* preservation status were recorded, and complete standard autopsies were performed.

The PMP-BS database was queried for all cetaceans stranded between February/16 and November/18. As inclusion criteria, only animals in ‘*fresh*’ (code 2) or moderate (code 3) *postmortem* condition (*16*) with sufficient formalin-fixed paraffin-embedded (FFPE) tissue and frozen tissue (−20°C) for immunohistochemical and molecular analysis, respectively, were included.

### Histopathology and immunohistochemistry

The histopathologic and immunohistochemical evaluation were performed at

LAP-UEL. Samples of the central nervous system (CNS), lungs and lymph nodes were collected, immersed in a 10% buffered formalin solution, and routinely processed for histopathologic evaluation stained with hematoxylin-eosin (H&E). Immunohistochemistry (IHC) was performed as described (*17*) using a monoclonal IgG_2b_ (kappa light chain) antibody specific for Canine Distemper Virus (CDV) nucleoprotein (CDV-NP, VRMD, Inc.; Pullman, WA, USA) as the primary antibody (1:500 dilution) for CNS, lung, and lymph nodes samples; and a polyclonal IgG anti-*Toxoplasma gondii* antibody (LS-C312239, LifeSpan BioSciences, Inc.; Seattle, WA, USA) as the primary antibody (1:200 dilution) for CNS sample. IHC anti-*T. gondii* was performed only on animals with typical histopathologic lesions of toxoplasmosis. Positive and negative controls were included in each assay. Positive antigen controls consisted of FFPE canine and feline brain tissue sections known to be positive for CDV (*17*) and *Toxoplasma gondii* antigens, respectively; negative controls consisted of the diluents of the primary antibody which substituted each primary antibody.

### Molecular analysis

Lungs and CNS samples were collected and stored at −20°C at LEC-UFPR. The CNS samples consisted predominantly of cerebellum, however, in a few cases a pool of cerebellum and cortex samples was performed. The samples were transported to LV-UEL frozen on dry ice in thermal transport boxes where molecular assays were performed.

Suspension (10% w/v) of tissue samples were prepared from a 100 mg portion of each sample and mechanically disrupted (MagNa Lyser Instrument, Roche Diagnostics, Mannheim, Germany), homogenized in 0.01 M phosphate buffered saline at pH 7.2, and clarified by centrifugation at 2,000 × g for 10 minutes. The nucleic acid extraction was fulfilled with 250 μL suspension aliquots pre-treated with proteinase K, using a combination of phenol/chloroform/isoamyl alcohol and silica/guanidine isothiocyanate methods (*18, 19*). The reverse transcription polymerase chain reaction (RT-PCR) was carried out using CeMV-specific primers for a 374-bp conserved fragment of the phosphoprotein (P) previously described (*14*) with modifications: 5’-ATGTTTATGATCAC**G**GCGGT-3’ (forward) and 5’-T**C**GGGTTGCACCACTTGTC-3’ (reverse) (bold letters indicating the modifications). Amplification reactions were performed using 5 μL of the extracted nuclei acid in a thermocycler (Swift™ MaxPro Thermal Cycler, Esco Healthcare Pte, Singapore). Negative control consisted of sterile ultrapure water, that was used in all the nucleic acid extractions and subsequent procedures; and positive controls were kindly provided by Catão-Dias JL and Groch KR and consisted of cetacean spleen, lungs, and brain tissue fragments known to be positive in RT-PCR for CeMV. The amplified products were analyzed by electrophoresis on a 2% agarose gel in TBE buffer, pH 8.4 (89 mM Tris; 89 mM boric acid; 2 mM EDTA), stained with ethidium bromide (0.5 μg/mL) and visualized under UV light.

## Results

### Animals

A total of 325 records of autopsied stranded cetaceans were found between February 2016 and November 2018. Of these, a total of 40 animals met the adopted criteria and *Sotalia guianensis* (SG, 30/40, 75%) was the most common species, while other species such as *Pontoporia blainvillei* (PB, 4/40, 10%), *Tursiops truncatus* (TT, 2/40, 5%), *Stenella* spp. (S, 1/40, 2.5%), *Stenella frontalis* (SF, 1/40, 2.5%), *Balaenoptera acutorostrata* (BA, 1/40, 2.5%), and *Steno bredanensis* (SB, 1/40, 2.5%) were less frequently observed.

These animals were classified predominantly as code 3 (28/40, 70%) – moderate *postmortem* condition – and the remaining were classified as code 2 (12/40, 30%) – ‘fresh’ *postmortem* condition. Additionally, 23 (57.5%) animals were identified as male, 16 (40%) as female, and in one (2.5%) animal it was not possible to identify the sex; 21 (52%) were adult animals, 15 (38%) juvenile, and four (10%) calves. At gross examination, 27 (67.5%) animals presented a good body condition, 10 (25%) animals a poor body score, one (2.5%) was cachectic and two (5%) were undetermined. The most common anthropic interaction observed was bycatch (21/40, 52.5%) (Figure 1), the absence of any suggestive lesions of anthropic interaction was revealed in seven animals (17.5%), and one animal (2.5%) presented solid waste interaction. It was not possible to establish whether there was anthropic interaction in 11 (27.5%) animals due to skin sloughing as result of *postmortem* changes. The individual stranding date, species, *postmortem* condition, gender, stage of development, body score, stranding condition, and anthropic interaction are summarized in appendix A.

**Figure 1.**
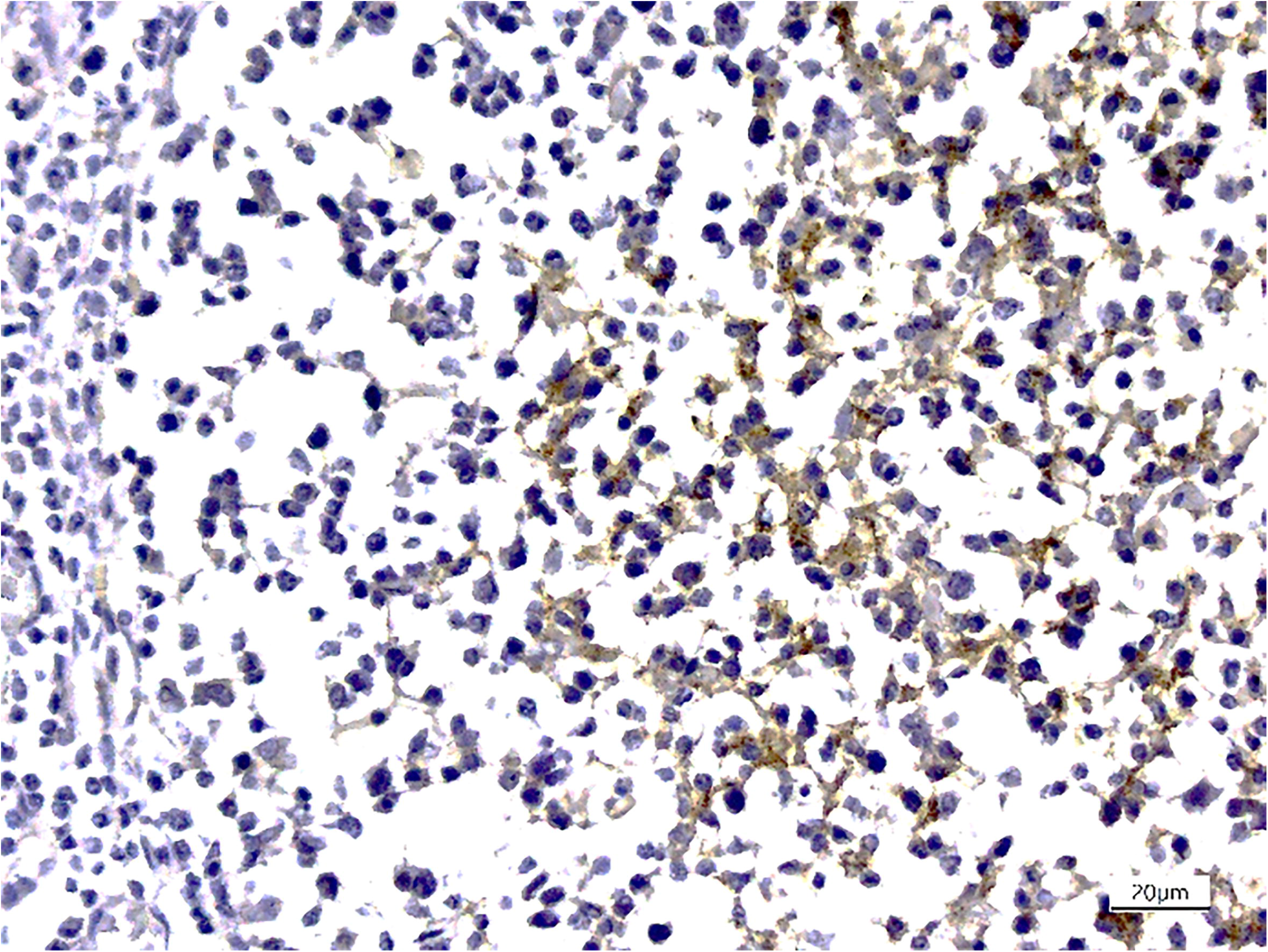
An franciscana dolphin calf (*Pontoporia blainvillei*) exhibiting a gillnet attached to the rostrum, compatible with bycatch.

### Histopathology and Immunohistochemistry

The main lungs histopathologic findings observed were granulomatous bronchopneumonia (9/40, 22.5%), lymphoplasmacytic interstitial pneumonia (8/40, 20%) (Figure 2a, 3a), and eosinophilic and granulomatous bronchopneumonia (7/40, 17.5%); whereas suppurative bronchopneumonia (2/40, 5%) and pyogranulomatous pneumonia (2/40, 5%) were less commonly observed. Interestingly, it was observed different patterns of pneumonia in 5 animals (5/40, 12.5%), in which two (5%) of them had lymphoplasmacytic interstitial pneumonia and suppurative bronchopneumonia; and three (7.5%) animals had eosinophilic and granulomatous bronchopneumonia and lymphoplasmacytic interstitial pneumonia, concomitantly. Pulmonary nematodes (*Halocercus brasiliensis*) were observed in eight (8/40, 20%) animals, intralesional bacteria in four (4/40, 10%) animals and the involvement by two different etiological agents also was observed in three animals (3/40, 7.5%) - two animals affected by bacteria and *H. brasiliensis*, and one by bacteria and dichotomous (45 degrees) branching, fungus hyphae with frequent separation suggestive of *Aspergillus* spp. (Figure 3b). Non-specific pulmonary lesions such as pulmonary edema (16/40, 40%) and passive congestion (2/40, 5%) were also observed. Formation of syncytial cells and inclusion bodies were rarely observed. Lymphoid depletion was the lesion most frequently observed in lymph nodes (Figure 3e) (16/40, 40%), and multifocal areas of necrosis (10/40, 25%) associated or not with lymphoid depletion.

**Figure 2.**
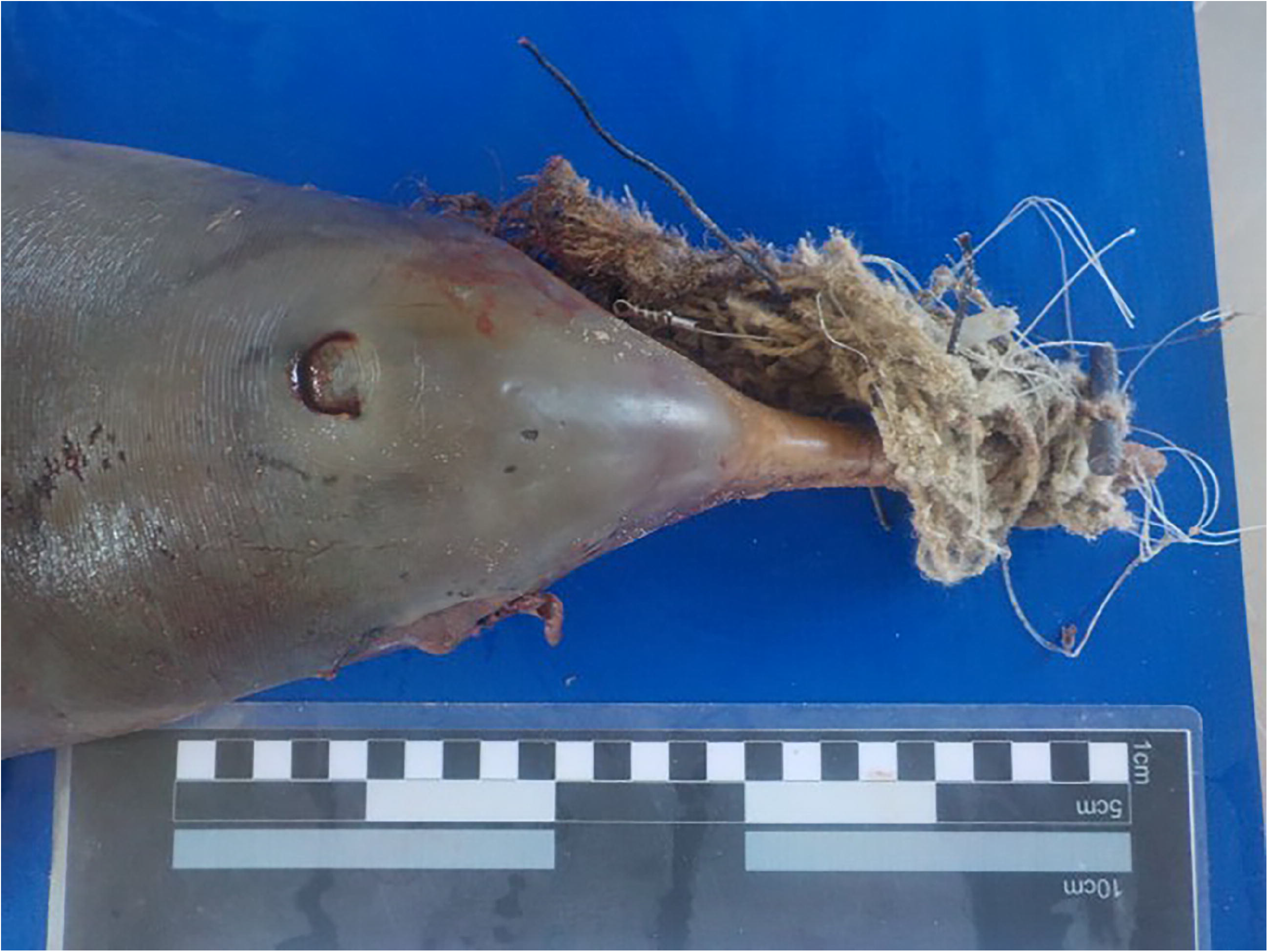
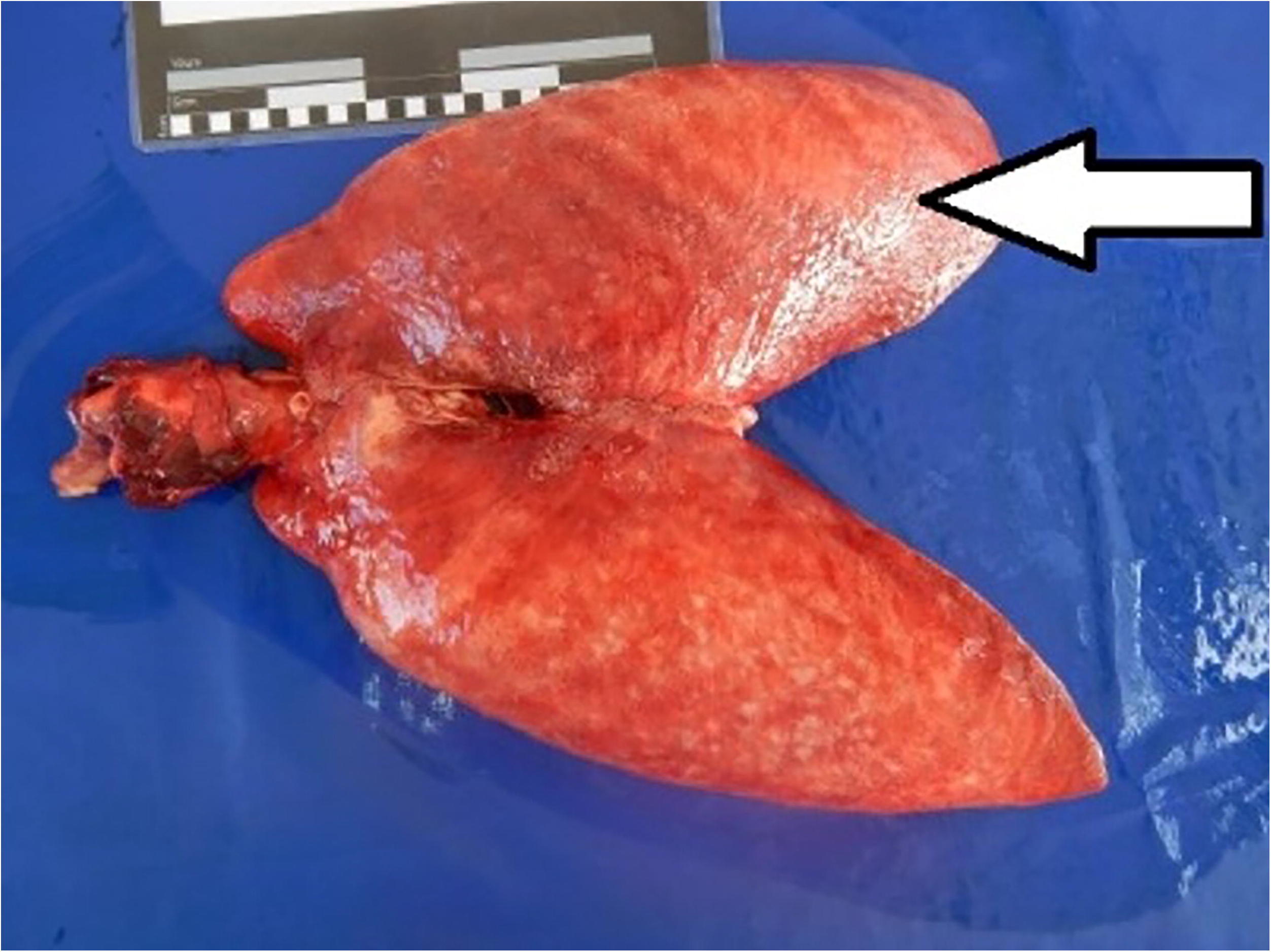
a) Uncollapsed and expanded lungs showing rib prints suggesting diffuse interstitial pneumonia (arrow). b) Multifocal to coalescing reddish areas suggesting encephalitis. Brain, *Sotalia guianensis*.

**Figure 3.**
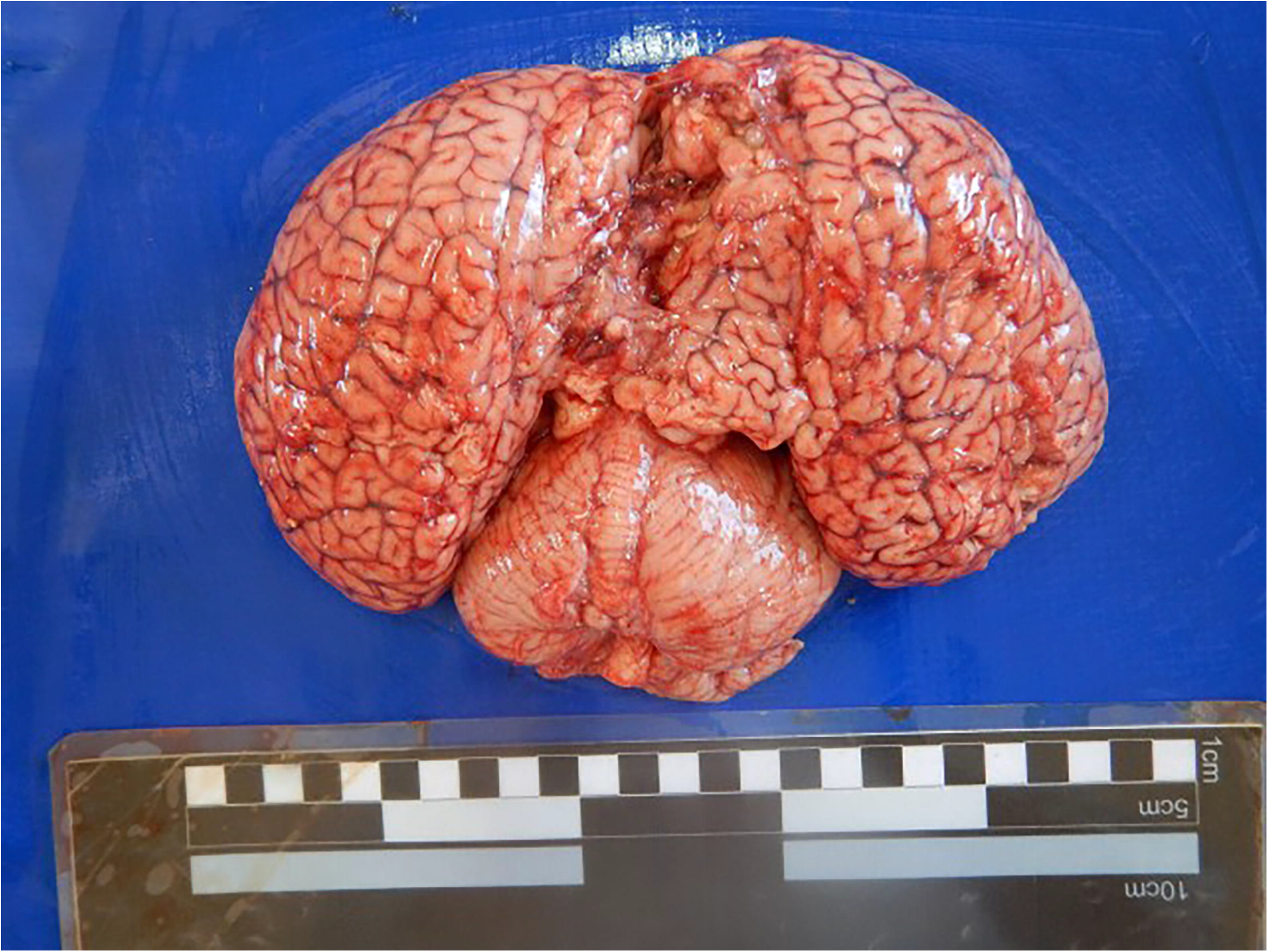
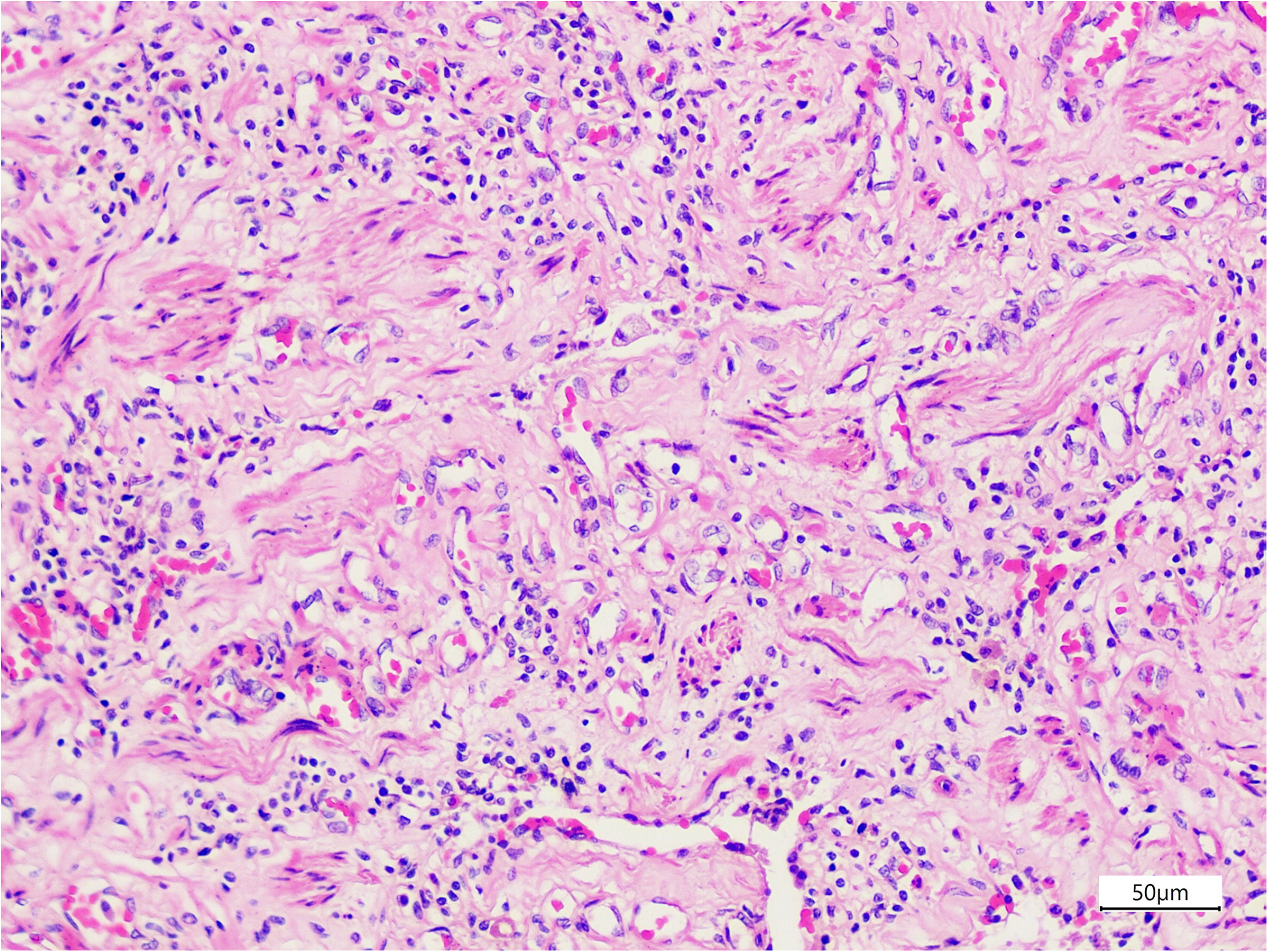
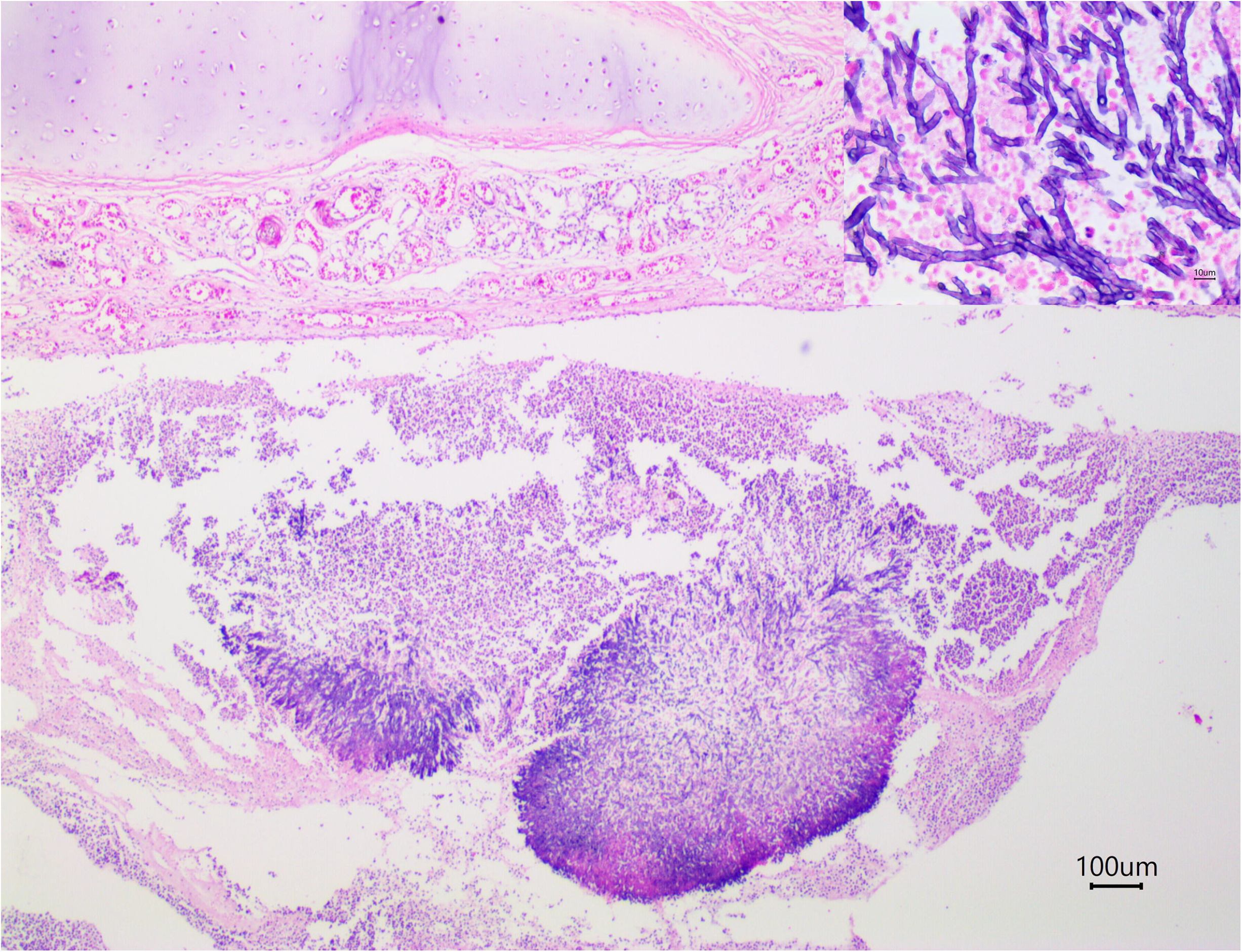
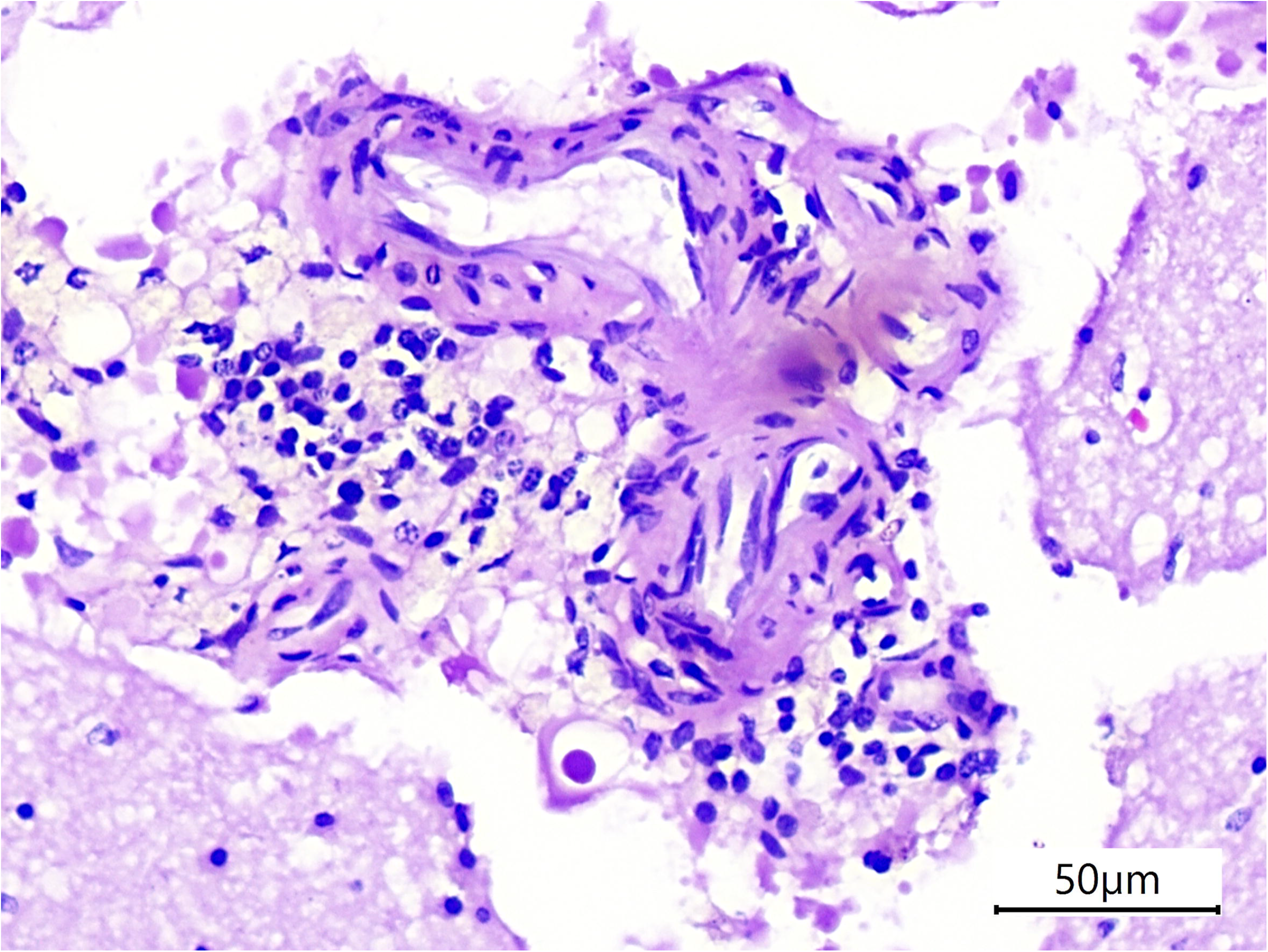
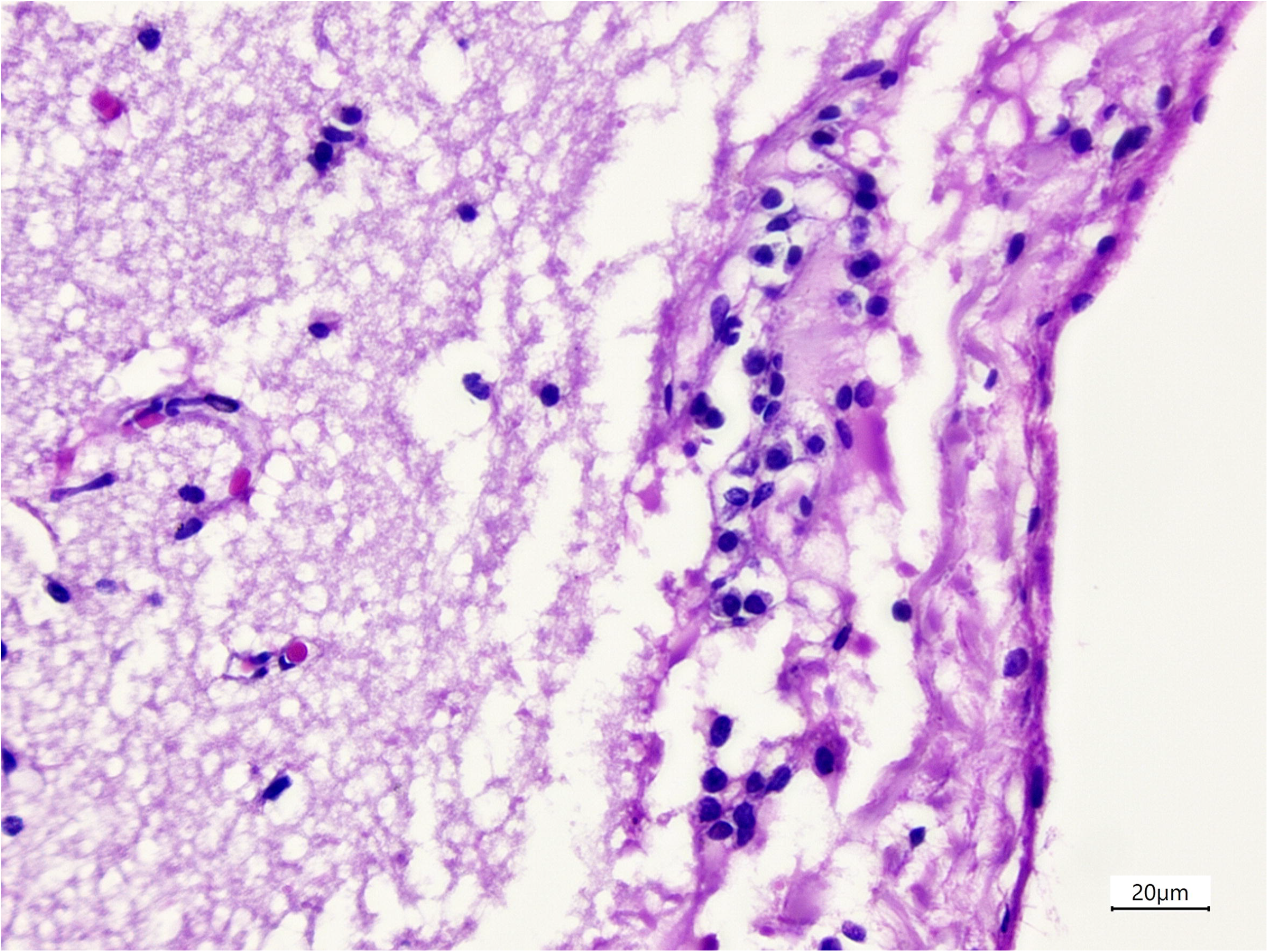
a) Lymphoplasmacytic inflammation in the pulmonary interstitium and atelectasis with random scattered areas of fibrosis. Lung, *S. guianensis* (H.E., 200x, bar: 50 μm). b) Fungal hyphae mixed with predominantly neutrophilic inflammatory infiltrate, cellular debris and red blood cells in the bronchus. Lung, *S. guianensis* (H.E., 40x, bar: 100 μm). Inset upper right: Septate, dichotomous (45°) branching hyphae suggestive of *Aspergillus* spp. c) Perivascular lymphoplasmacytic cuffing in cerebral cortex. Brain, *S. guianensis* (H.E., 200x, bar: 50 μm). d) Lymphocytic inflammation in cortical leptomeninges. Brain, *S. guianensis* (H.E., 400x, bar: 20 μm). e) Diffuse rarefaction of lymphoid cells with multifocal areas of necrosis with central mineralization (*). Pulmonar lymph node, *S. guianensis* (H.E., 40x, bar: 100 μm).

The CNS lesions were less frequent and included non-suppurative encephalitis (1/40, 2.5%) (Figure 2b, 3c), non-suppurative encephalitis with multifocal gliosis and intralesional protozoa (1/40, 2.5%), non-suppurative meningitis with neuronal necrosis and satellitosis (1/40, 2.5%) (Figure 3d), non-suppurative meningoencephalomyelitis with gliosis and intralesional protozoa (1/40, 2.5%), satellitosis (1/40, 2.5%), and polioencephalomalacia (1/40, 2.5%). These lesions were all multifocal, mild, and observed predominantly in cerebral cortex; the cerebellum was less frequently affected. The mild perivascular cuffs ranged from one up to two layers. Additionally, protozoa cysts were observed surrounded by areas of gliosis in the white matter of two animals with non-suppurative encephalitis and meningoencephalomyelitis (animals #11 and #18); and parasites eggs of *Nasitrema globicephalae* were observed at cerebellum white matter in one animal. Histopathologic evaluation also revealed non-specific lesions such as encephalic hemorrhage (5/40, 12.5%) and encephalic congestion (6/40, 15%). All histopathologic data is summarized at appendix B.

At immunohistochemical evaluation, a total of 11 (11/40, 27.5%) animals presented positive immunoreactivity anti-CeMV antigens in FFPE lung (Figure 4a, 4b) and/or CNS sections (Figure 4c). Of those, we found 10 (10/40, 90.9%) animals with CeMV antigens exclusively in lungs sections associated with pneumonia, except for animal #9 which did not present pulmonary inflammatory lesions; and one (1/40, 9.1%) animal presented positive immunoreactivity in both organs, associated with lymphohistiocytic interstitial pneumonia and non-suppurative meningitis, satellitosis and neuronal necrosis. All animals which presented intracytoplasmic immunoreactivity in lungs or CNS also presented intracytoplasmic immunoreactivity within lymphocytes and macrophages in lymph nodes (Figure 4e). Additionally, the intralesional protozoa observed in animals #11 and #18 at histopathologic evaluation were positively immunoreactive in anti-*Toxoplasma gondii* IHC (Figure 4d). All immunohistochemical data are summarized at appendix B.

**Figure 4.**
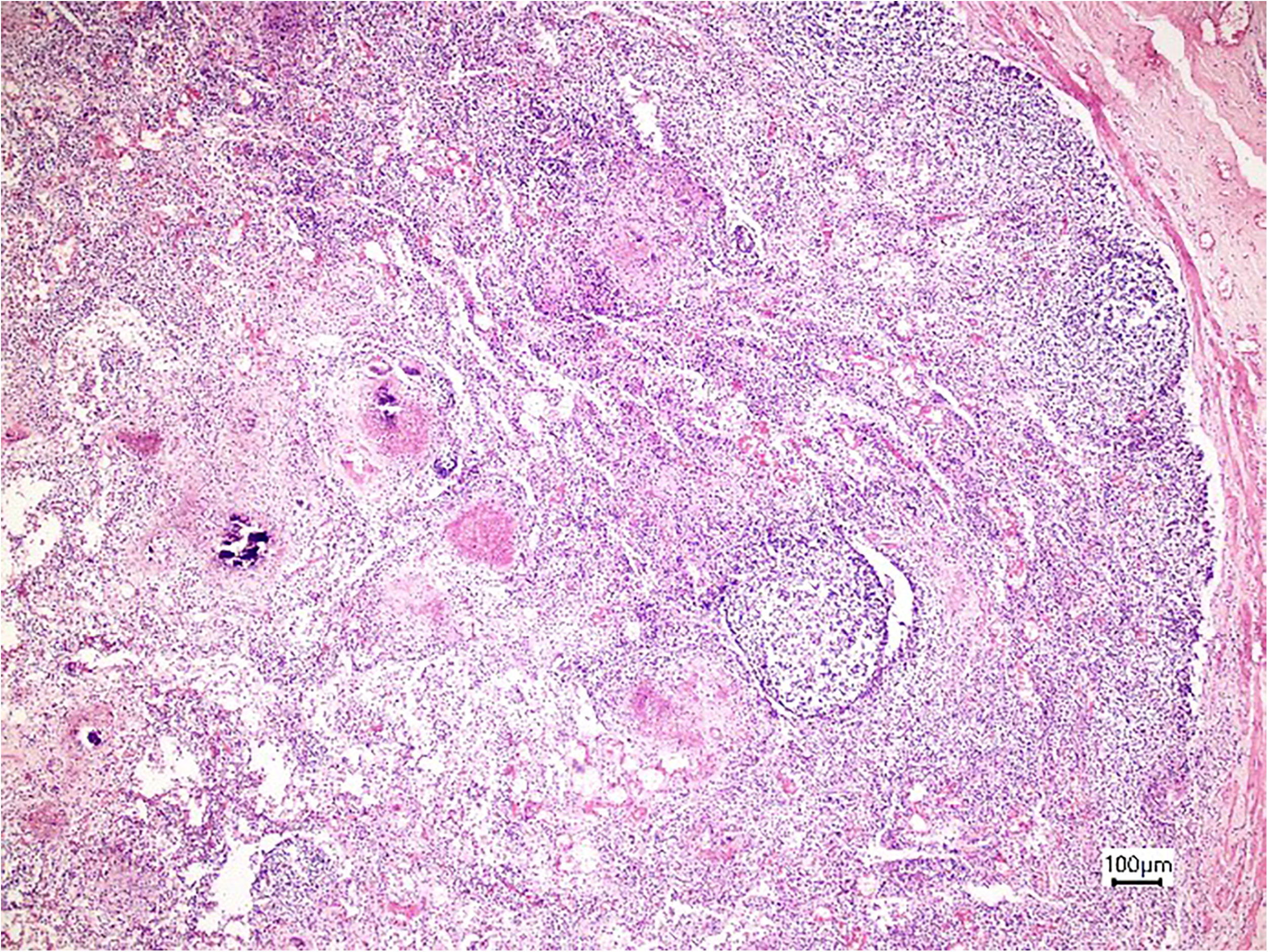
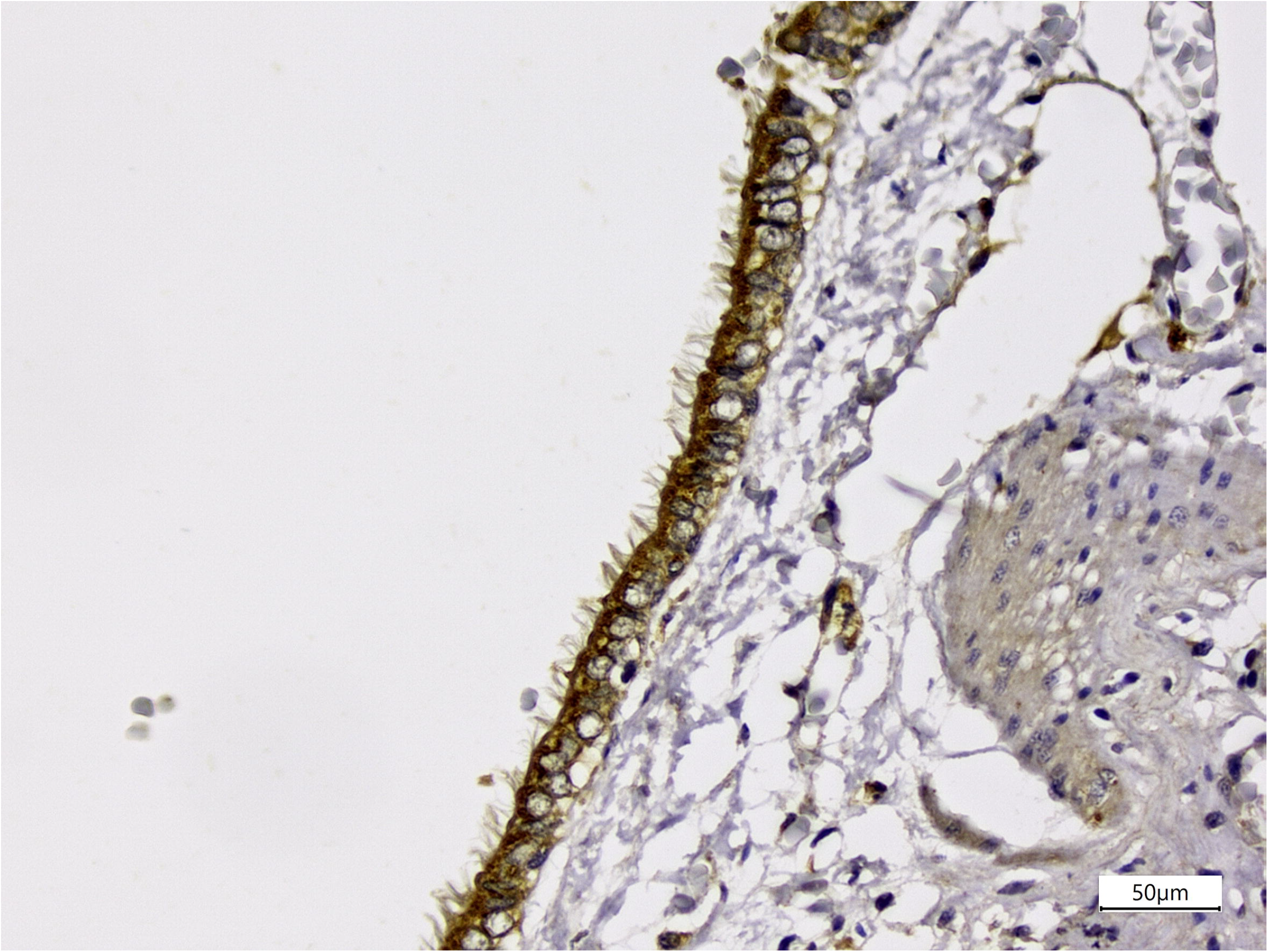
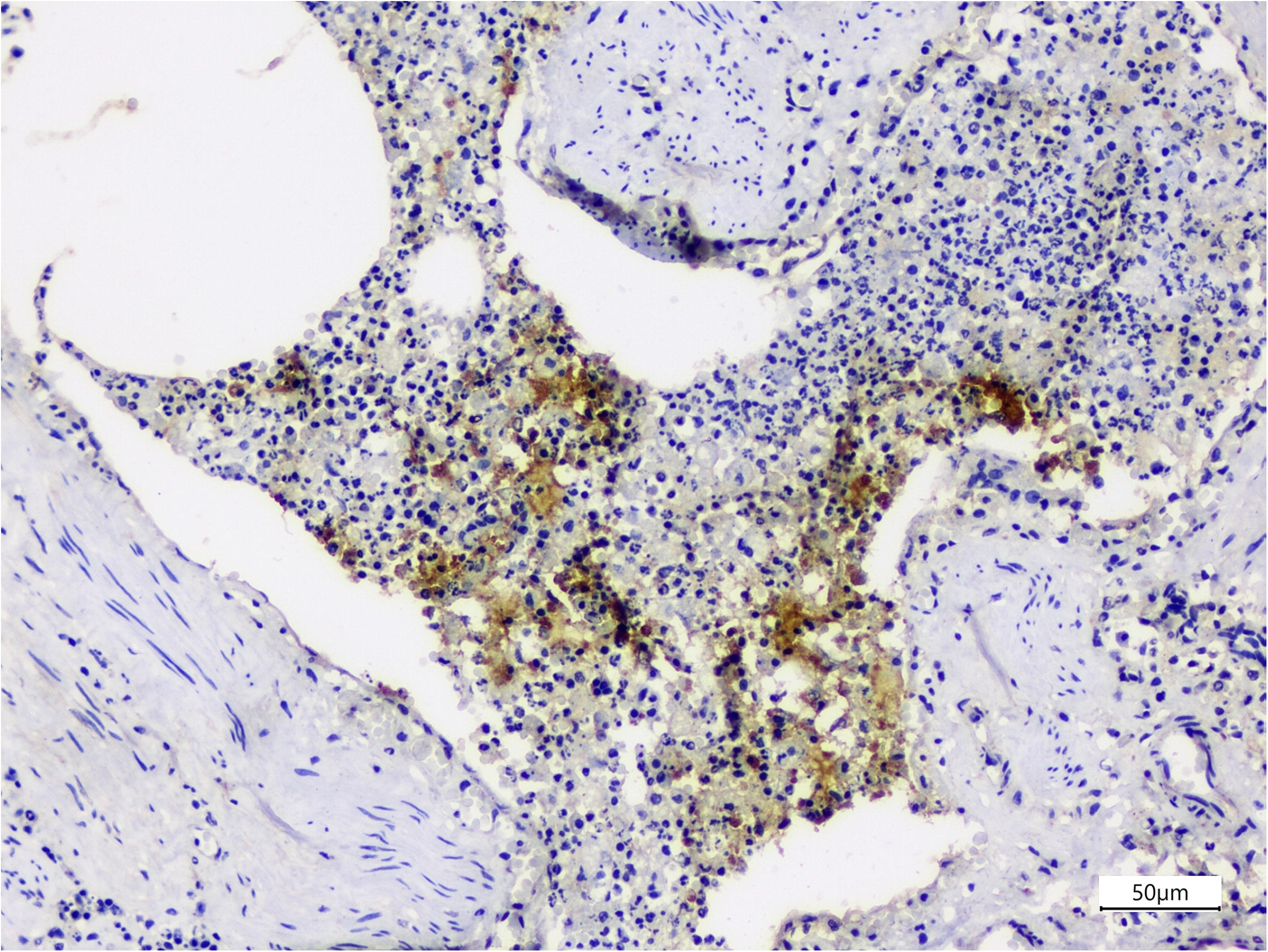
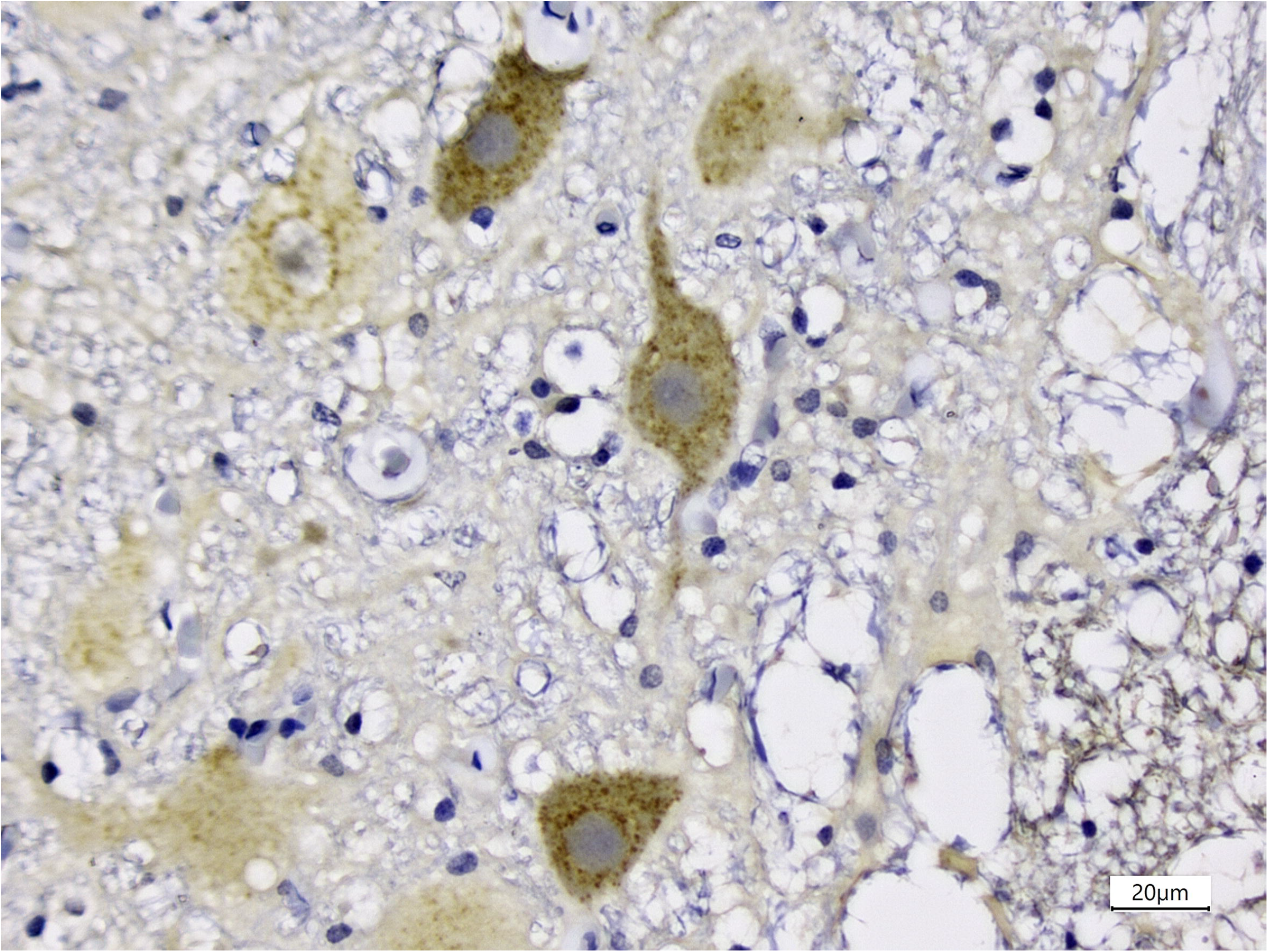
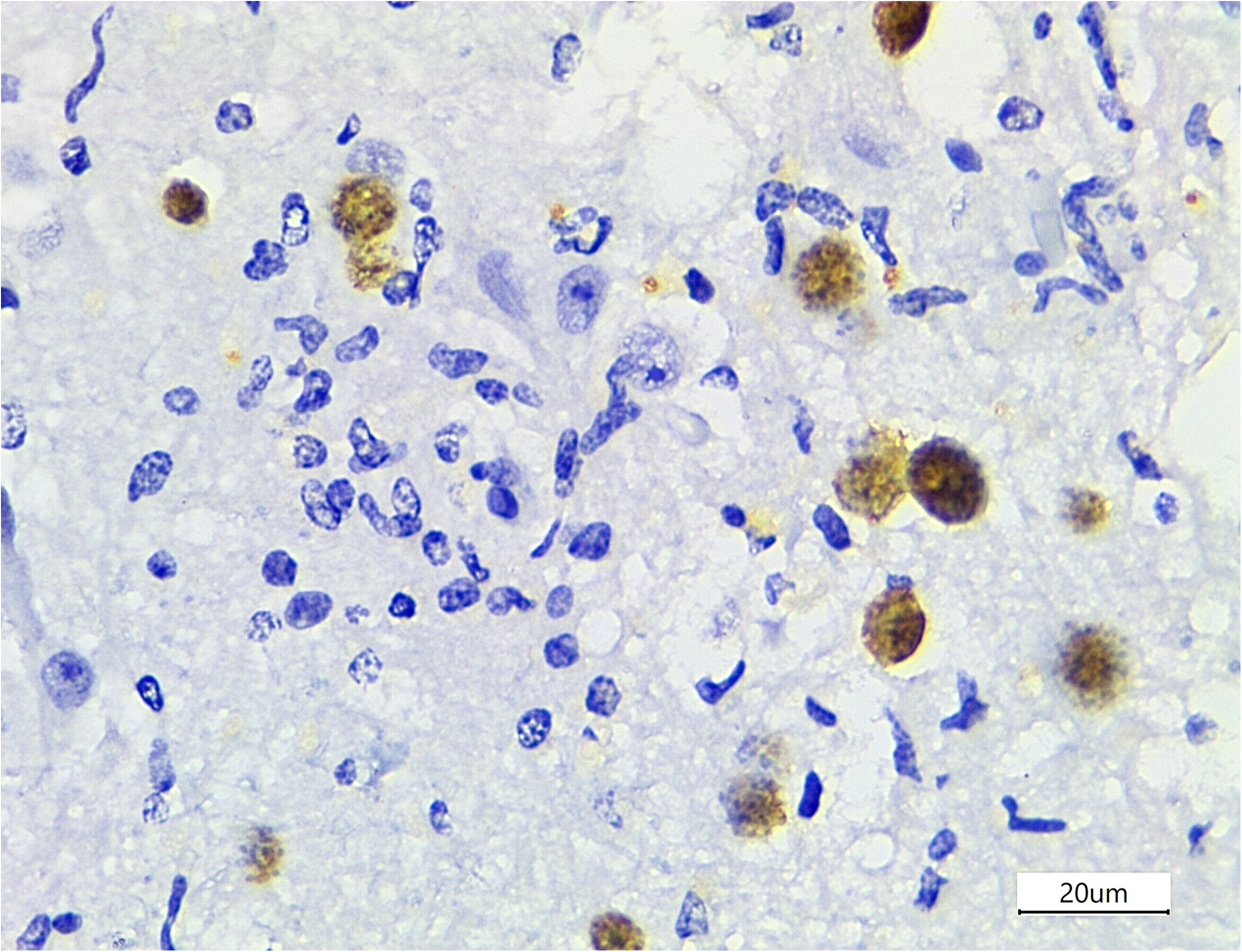
a) Consistent intracytoplasmic immunolabeling anti-CeMV within bronchial epithelial cells. Lung, *S. guianensis* (IHC anti-CeMV, 200x, bar: 50 μm). b) Mixed intrabronchiolar inflammation with consistent intracytoplasmic immunolabeling within lymphocytes and macrophages. Lung, *S. guianensis* (IHC anti-CeMV, 200x, bar: 50 μm). c) Consistent intracytoplasmic immunolabeling within spinal cord neurons. Spinal cord, *S. guianensis* (IHC anti-CeMV, 200x, bar: 20 μm). d) Consistent intracytoplasmic immunolabeling within protozoa cysts around a focal area of gliosis. Cerebral cortex, *S. guianensis* (IHC anti-*Toxoplasma gondii*, 600x, bar: 20 μm). e) Consistent intracytoplasmic immunolabeling anti-CeMV antigens within lymphocytes in a depleted lymph node. Pulmonary lymph node, *S. guianensis* (IHC anti-CeMV, 400x, bar: 20 μm).

## Discussion

The marine mammals research area is challenging since there are many uncontrollable factors such as the time between the death in the sea, the stranding and the autopsy examination leading to variable *postmortem* changes; environmental adversities that would difficult the beach monitoring; and geographical conditions which do not allow the daily or weekly monitoring in a few areas. Although the factors mentioned above may difficult research, we verified the occurrence of CeMV under non-epizootic conditions at a world heritage in Southern Brazil in cetaceans along off the Paraná coast that died between 2016-2018 due to a combination of pathologic and immunohistochemical findings.

Due to the fact that PMV and DMV are antigenically closely related, demonstrating a similar reaction pattern in indirect ELISA assay with monoclonal antibodies raised against CDV, PDV, PPRV (*peste de petits ruminants*) and RPV (rinderpest virus) proteins (*20, 21*), and that antigenic similarity between different strains of morbillivirus were also observed in serological studies (*22, 23*), we used a monoclonal antibody anti-CDV nucleoprotein as primary antibody for cetacean morbillivirus researching. Although it was not possible to identify the virus in RT-PCR assay, the term was used ‘CeMV’ because it is the most likely virus in this species.

The difference between the number of records of autopsied cetaceans and the number of animals evaluated was primarily due to advanced *postmortem* conditions and lack of tissue samples for histopathologic and/or molecular analysis in a few cases. The most common species herein described was *Sotalia guianensis,* probably because of its near-shore distribution and site fidelity (*24*), which makes it more susceptible to the effects of anthropogenic activities (*25*). Other species such as *Tursiops truncatus, Stenella frontalis, Balaenoptera acutorostrata*, and *Steno bredanensis* were less frequent, similarly to other study carried out in the same region (*26*).

Most animals were in a good body condition and suggestive lesions of bycatch were observed. Of 21 cetaceans with variable degrees of bycatch lesions, only two animals were not in good body condition; one was in a poor body condition and the other could not be determined as result of scavenger animals. Historically, *S. guianensis* and *P. blainvillei* are the most common and regionally caught cetaceans (*27, 28*), maintaining a high vulnerability off Paraná (*26, 29*). The frequency of bycatch lesions in cetaceans described herein corroborates with previous study conducted in the same region (28/46, 61%) (*26*), and in southeastern (60/79, 76%) of Brazil (*30*). Additionally, considering that it was not possible to establish whether there was anthropic interaction in 11 animals, the frequency of bycatch may be higher, representing a serious threat to cetaceans’ population with important demographic consequences (*31*) and dramatical changes in the structure and function of marine ecosystems (*32–34*). Consistent intracytoplasmic immunolabeling anti-CeMV was found in five (5/21, 23.8%) of those animals with bycatch lesions, reinforcing that anthropogenic activities are a leading cause of cetacean stranding in Paraná, but underlying pre-existing diseases might contribute towards deaths also (*26*).

The main lungs histopathologic findings in animals infected by CeMV were lymphoplasmacytic interstitial pneumonia, and antigens anti-CeMV were found predominantly in epithelial cells and rarely in lymphocytes and macrophages surrounding the bronchus and bronchioles. The lymphocytic infiltrate associated with CeMV infected animals were predominantly peribronchial and/or peribronchiolar corroborating with a previous study (*22*), however, hyperplasia of bronchiolar epithelium, proliferation of type II pneumocytes and formation of syncytia were not a prominent feature. Two of these animals also presented different patterns of pneumonia; one with eosinophilic granulomatous bronchopneumonia and the other with suppurative pneumonia. The remaining animals presented granulomatous or suppurative inflammation. *Halocercus brasiliensis* was found in four animals with consistent immunolabeling anti-CeMV in IHC.

The most common lungs findings in cetaceans affected by CeMV are interstitial pneumonia (*1, 35-37*), however, the typical lesions of a CeMV infection could be considerably overlapped by secondary infectious agents, such as *Halocercus brasiliensis* (*8*), mainly in sub-acute or chronic infection as a consequence of severe immunosuppression (*38*). Although we did not observe typical histopathologic changes of CeMV and absence of secondary infectious agents in one animal with consistent intracytoplasmic immunolabeling anti-CeMV antigens within bronchial epithelial cells; lymphocytes in the lymph nodes parenchyma (pulmonary, mediastinal, cervical, submandibular, and prescapular); and endothelial cells, we strongly suspect of a systemic acute early developmental stage of disease.

Histopathologic changes in CNS were less frequently observed and comprised mainly of non-suppurative meningoencephalitis or encephalitis. Additionally, two animals with non-suppurative meningo and/or encephalitis with intralesional protozoa at histopathologic evaluation, had consistent immunolabeling anti-*Toxoplasma gondii* in cerebral cortex and cerebellum. Toxoplasmosis was often associated with immunosuppression followed by a morbillivirus infection (*35-37, 39*), however, CeMV antigens were not found at IHC in both animals. One animal presented mild typical CNS lesions of a CeMV infection, such as non-suppurative encephalitis, satelitosis, and neuronal necrosis (*2, 36, 37*). In IHC, consistent intracytoplasmic immunolabelling anti-CeMV antigens were found within neurons, glial cells, and rarely in endothelial cells.

In the present study, we observed CeMV antigens in 25% SG (10/40), however, only one SG presented CeMV antigens in both lungs and CNS samples with mild inflammatory changes in CNS histopathology. A comparative study revealed that the lack of neurodegenerative, reactive, or inflammatory changes in SG suggests low neurovirulence of the strain and that CNS involvement may occurs at a later stage of infection and that these animals succumbed too early for CNS pathology to be observed (*40*). Thus, it would explain why CNS lesions were mild and rarely observed. We did not have access to the whole brain to a wide and detailed sampling to map out the major affected regions in this species, and this should be considered as a limitation of the present study.

All samples were negative at RT-PCR analysis. We hypothesized that the negative results may have occurred due to (i) heterogenic distribution of the virus in organs, (ii) viral RNA degradation because of the poor preservation status frequently shown by tissue from stranded animals, or viral RNA degradation in the samples by RNases and/or temperature changes, (iii) low viral load in samples, (iv) different targets (gene P in RT-PCR and nucleoprotein in IHC) in the virus detection. Therefore, we strongly suggest the use of both RT-PCR and IHC when possible and store frozen samples at −80°C.

A previous study assessed the health of cetaceans in the same region and revealed the presence of CeMV antigens by IHC assay in five (8.7%) of the 57 animals between July 2007 and November 2012 (*26*). Four years later, we observed an increase of the prevalence of CeMV to 27% when evaluating animals from February 2016 to November 2018, approximately three times more in a shorter period than the previous study. These results may indicate a degradation of the environment and raise the concern about the conservation and preservation of cetaceans at Paraná’s coast, predominantly for SG species which was the first confirmed decline of a delphinid population from Brazilian waters (*24*), and it is a warning that marine environment requires protection and legal regulations for human activities.

In addition, emerging infectious diseases can alter populations abundances and cause major regime changes within communities (*41*), being frequently the result of the ecology of host and/or pathogen changes and are often driven by anthropogenic environmental modifications (*42–46*). The PEC has been severely impacted by anthropogenic activities such as port area (considered the most important handling site for grain and fertilizers in South America) (*47*); inorganic nutrient levels in untreated wastewater discharged from Paranaguá city by far exceeded legislated national threshold levels (*47*); and by shipwrecks, continuous and persistent oils spills, and trace-element contamination during traffic and unloading of organic matters (*48, 49*). Marine mammals are considered useful sentinels for emerging and reemerging infectious diseases and they are extremely sensitive to marine ecosystem health reflecting the magnitude and rapidity of environmental changes (*50*). In front of this, the systematic beach monitoring is an important and useful health assessment tool for a better understanding of anthropogenic activities consequences in marine environment and to drive mitigation and conservation actions to improve the ecosystem and human health.

In conclusion, diagnosis of CeMV is a challenge in areas in which epidemics episodes have not been recorded and due to postmortem changes. However, we observed a prevalence of 27.5% of CeMV with mainly pulmonary changes associated in stranded cetaceans off the coast of Paraná due to a combination of histological and immunohistochemical findings. This virus may play a major role in cetacean stranding and as well as acting as an immunosuppressive agent causing secondary infections of parasitic, bacterial and protozoa. The results herein described broadens the knowledge about CeMV in our region and it is the first study related to CeMV under natural circumstances in Brazil.

## Acknowledgments

The authors are grateful to Kátia R. Groch and José Luiz Catão Dias, who kindly provided samples used as positive control in molecular analysis. Victor Marutani and Ana Paula Bracarense acknowledge the fellowship from Conselho Nacional de Desenvolvimento Científico e Tecnológico – Brazil (CNPq) – Finance Code 132970/2018-0 and 308136/2018-7, respectively.

## Author Bio (first author only, unless there are only 2 authors)

Victor Hugo Brunaldi Marutani, DVM, is a postgraduate student at “Programa de pós-graduação em Ciência Animal – Universidade Estadual de Londrina” for master’s title. Interests are in infectious diseases in marine animals.

## Appendix A.

Biological data and stranding date on cetaceans stranded in Paraná coast.

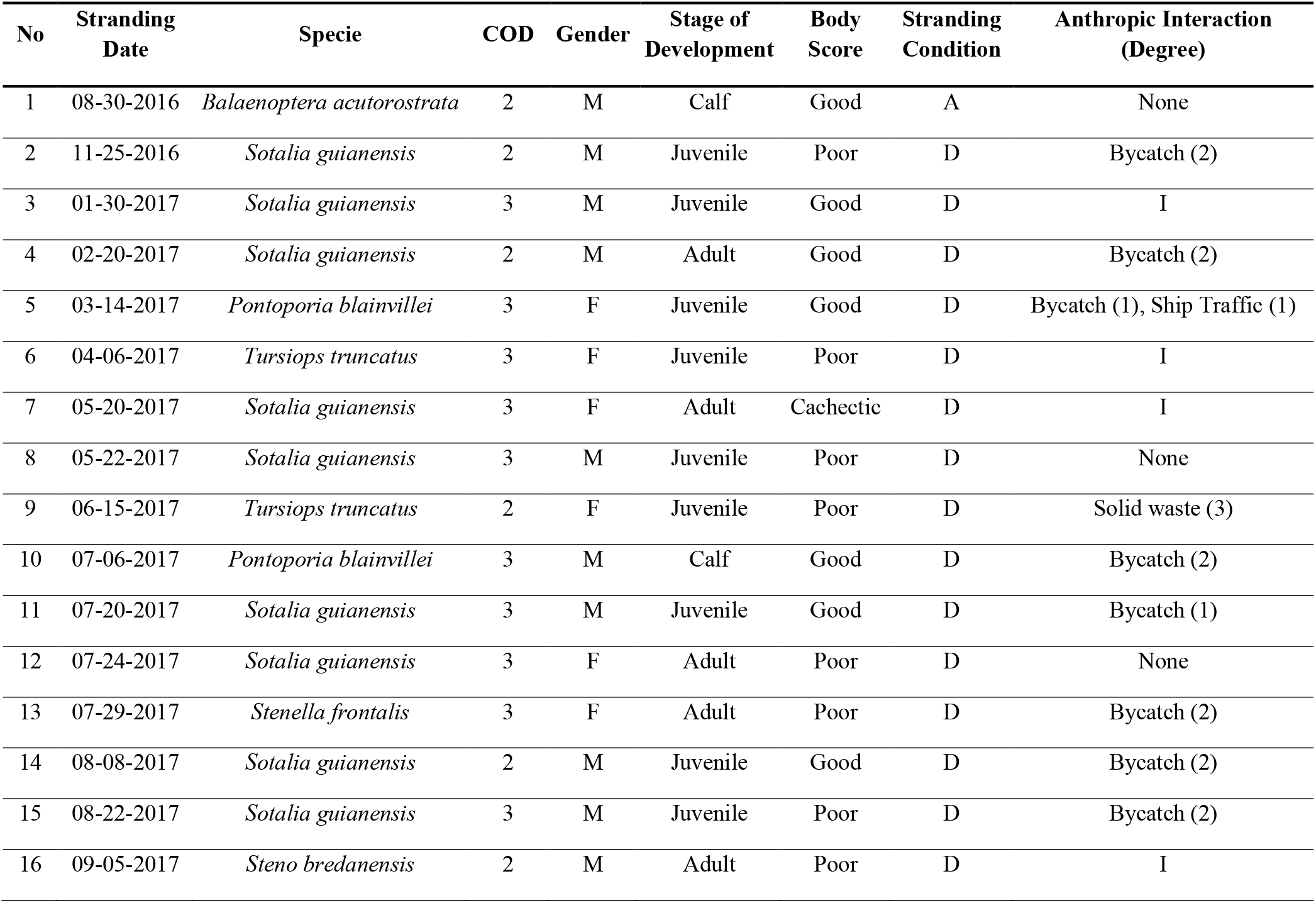
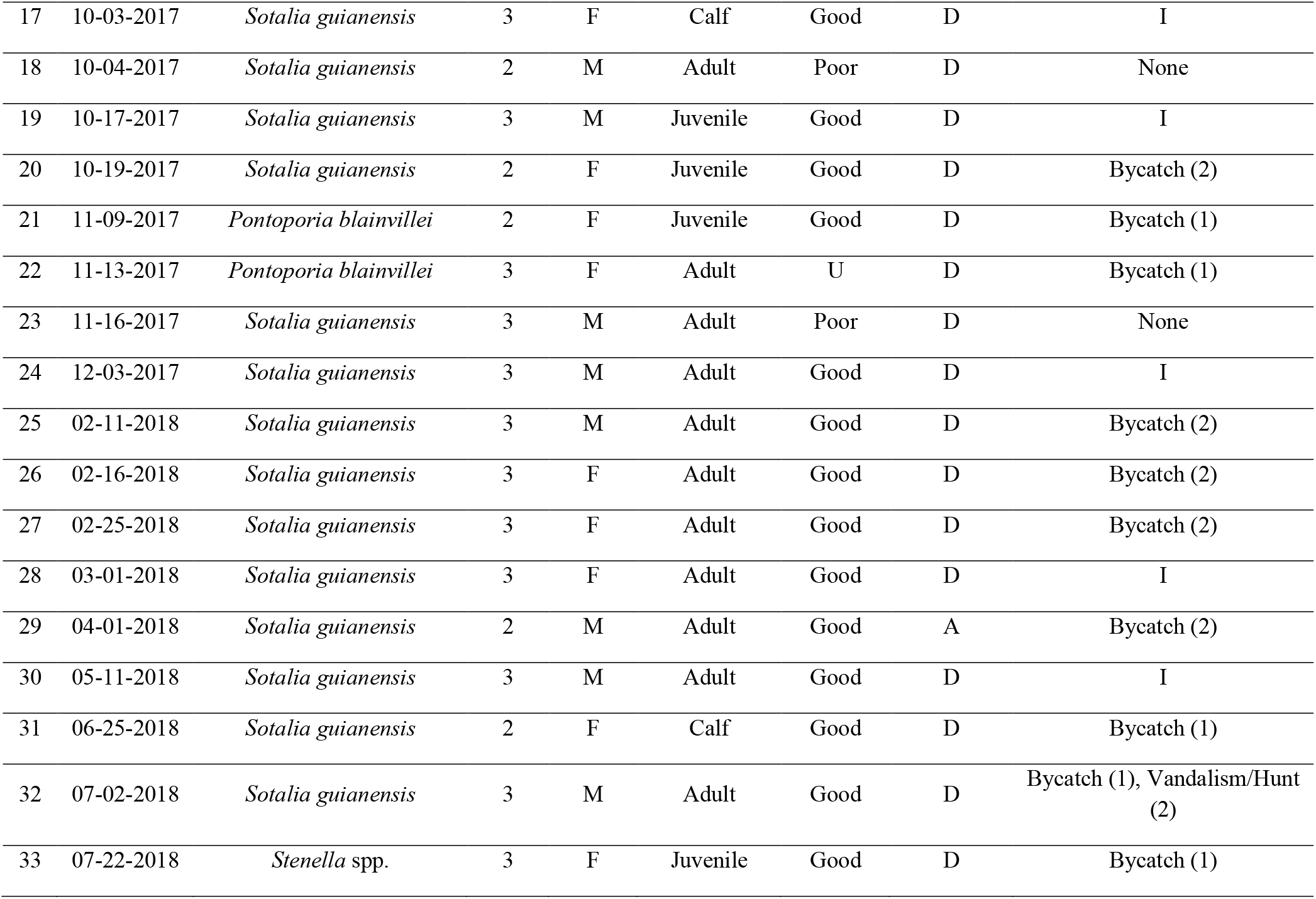
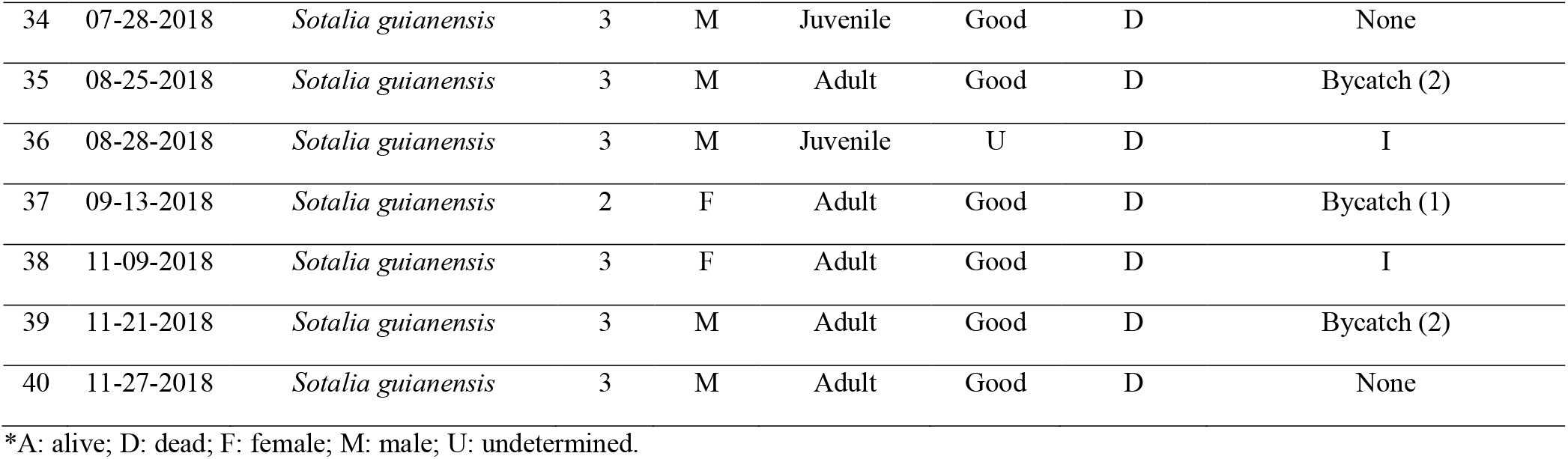

| No | Stranding Date | Specie | COD | Gender | Stage of Development | Body Score | Stranding Condition | Anthropic Interaction (Degree) |
| --- | --- | --- | --- | --- | --- | --- | --- | --- |
| 1 | 08-30-2016 | *Balaenoptera acutorostrata* | 2 | M | Calf | Good | A | None |
| 2 | 11-25-2016 | *Sotalia guianensis* | 2 | M | Juvenile | Poor | D | Bycatch (2) |
| 3 | 01-30-2017 | *Sotalia guianensis* | 3 | M | Juvenile | Good | D | I |
| 4 | 02-20-2017 | *Sotalia guianensis* | 2 | M | Adult | Good | D | Bycatch (2) |
| 5 | 03-14-2017 | *Pontoporia blainvillei* | 3 | F | Juvenile | Good | D | Bycatch (1), Ship Traffic (1) |
| 6 | 04-06-2017 | *Tursiops truncatus* | 3 | F | Juvenile | Poor | D | I |
| 7 | 05-20-2017 | *Sotalia guianensis* | 3 | F | Adult | Cachectic | D | I |
| 8 | 05-22-2017 | *Sotalia guianensis* | 3 | M | Juvenile | Poor | D | None |
| 9 | 06-15-2017 | *Tursiops truncatus* | 2 | F | Juvenile | Poor | D | Solid waste (3) |
| 10 | 07-06-2017 | *Pontoporia blainvillei* | 3 | M | Calf | Good | D | Bycatch (2) |
| 11 | 07-20-2017 | *Sotalia guianensis* | 3 | M | Juvenile | Good | D | Bycatch (1) |
| 12 | 07-24-2017 | *Sotalia guianensis* | 3 | F | Adult | Poor | D | None |
| 13 | 07-29-2017 | *Stenella frontalis* | 3 | F | Adult | Poor | D | Bycatch (2) |
| 14 | 08-08-2017 | *Sotalia guianensis* | 2 | M | Juvenile | Good | D | Bycatch (2) |
| 15 | 08-22-2017 | *Sotalia guianensis* | 3 | M | Juvenile | Poor | D | Bycatch (2) |
| 16 | 09-05-2017 | *Steno bredanensis* | 2 | M | Adult | Poor | D | I |
| 17 | 10-03-2017 | *Sotalia guianensis* | 3 | F | Calf | Good | D | I |
| 18 | 10-04-2017 | *Sotalia guianensis* | 2 | M | Adult | Poor | D | None |
| 19 | 10-17-2017 | *Sotalia guianensis* | 3 | M | Juvenile | Good | D | I |
| 20 | 10-19-2017 | *Sotalia guianensis* | 2 | F | Juvenile | Good | D | Bycatch (2) |
| 21 | 11-09-2017 | *Pontoporia blainvillei* | 2 | F | Juvenile | Good | D | Bycatch (1) |
| 22 | 11-13-2017 | *Pontoporia blainvillei* | 3 | F | Adult | U | D | Bycatch (1) |
| 23 | 11-16-2017 | *Sotalia guianensis* | 3 | M | Adult | Poor | D | None |
| 24 | 12-03-2017 | *Sotalia guianensis* | 3 | M | Adult | Good | D | I |
| 25 | 02-11-2018 | *Sotalia guianensis* | 3 | M | Adult | Good | D | Bycatch (2) |
| 26 | 02-16-2018 | *Sotalia guianensis* | 3 | F | Adult | Good | D | Bycatch (2) |
| 27 | 02-25-2018 | *Sotalia guianensis* | 3 | F | Adult | Good | D | Bycatch (2) |
| 28 | 03-01-2018 | *Sotalia guianensis* | 3 | F | Adult | Good | D | I |
| 29 | 04-01-2018 | *Sotalia guianensis* | 2 | M | Adult | Good | A | Bycatch (2) |
| 30 | 05-11-2018 | *Sotalia guianensis* | 3 | M | Adult | Good | D | I |
| 31 | 06-25-2018 | *Sotalia guianensis* | 2 | F | Calf | Good | D | Bycatch (1) |
| 32 | 07-02-2018 | *Sotalia guianensis* | 3 | M | Adult | Good | D | Bycatch (1), Vandalism/Hunt<br>(2) |
| 33 | 07-22-2018 | *Stenella* spp. | 3 | F | Juvenile | Good | D | Bycatch (1) |
| 34 | 07-28-2018 | *Sotalia guianensis* | 3 | M | Juvenile | Good | D | None |
| 35 | 08-25-2018 | *Sotalia guianensis* | 3 | M | Adult | Good | D | Bycatch (2) |
| 36 | 08-28-2018 | *Sotalia guianensis* | 3 | M | Juvenile | U | D | I |
| 37 | 09-13-2018 | *Sotalia guianensis* | 2 | F | Adult | Good | D | Bycatch (1) |
| 38 | 11-09-2018 | *Sotalia guianensis* | 3 | F | Adult | Good | D | I |
| 39 | 11-21-2018 | *Sotalia guianensis* | 3 | M | Adult | Good | D | Bycatch (2) |
| 40 | 11-27-2018 | *Sotalia guianensis* | 3 | M | Adult | Good | D | None |

\*A: alive; D: dead; F: female; M: male; U: undetermined.

## Appendix B.

Summarized data of histopathologic and immunohistochemical findings.

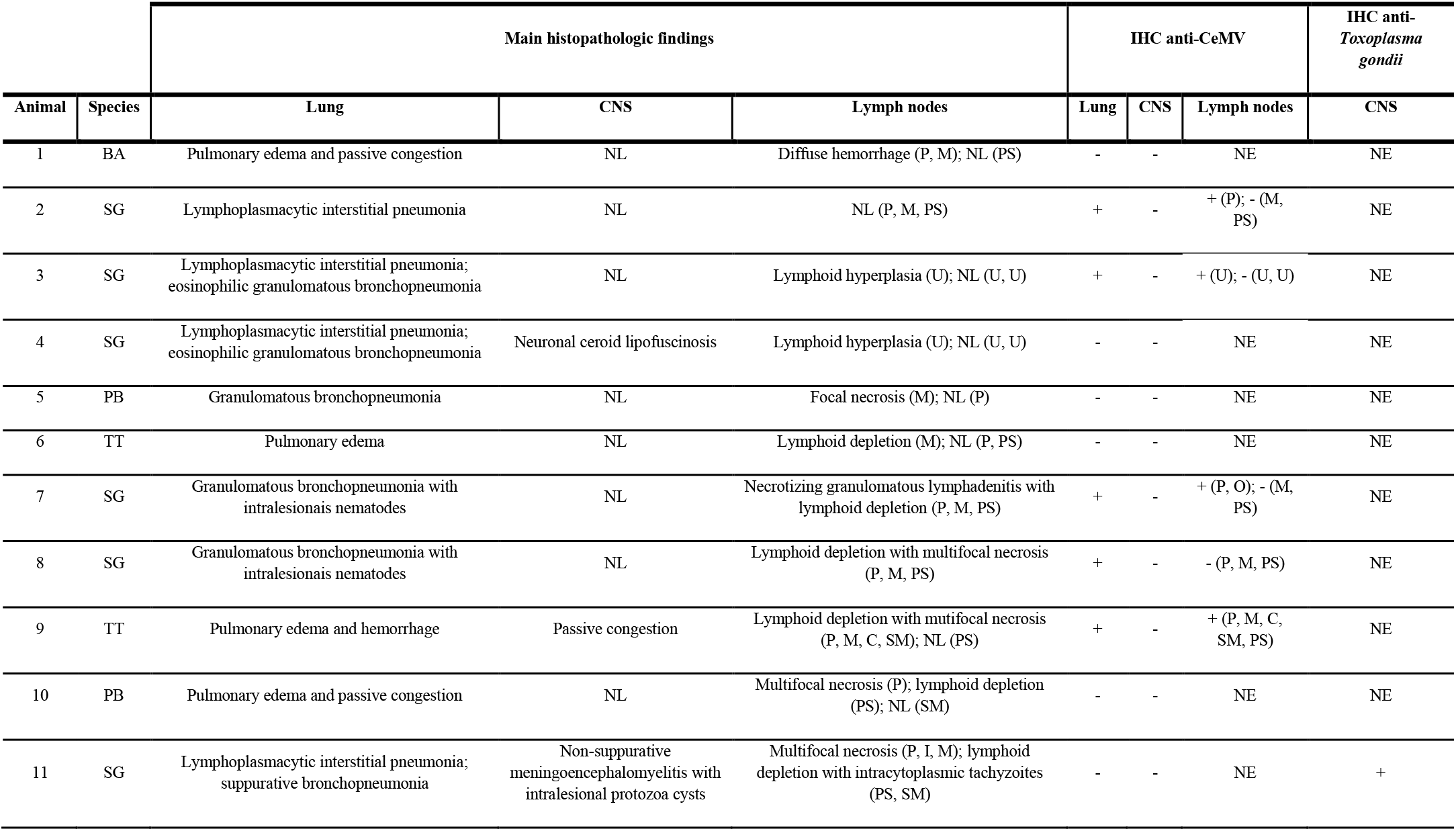
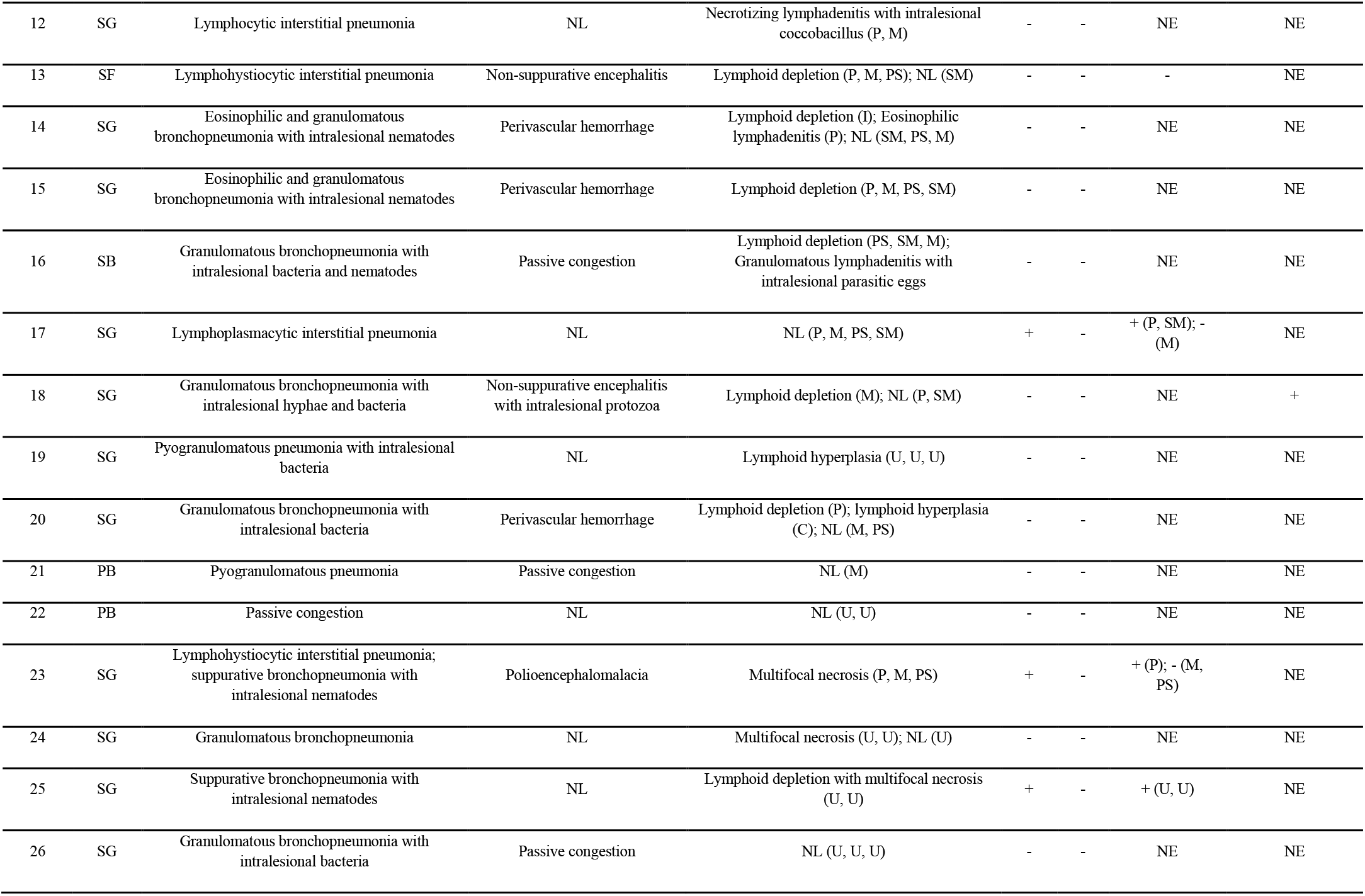
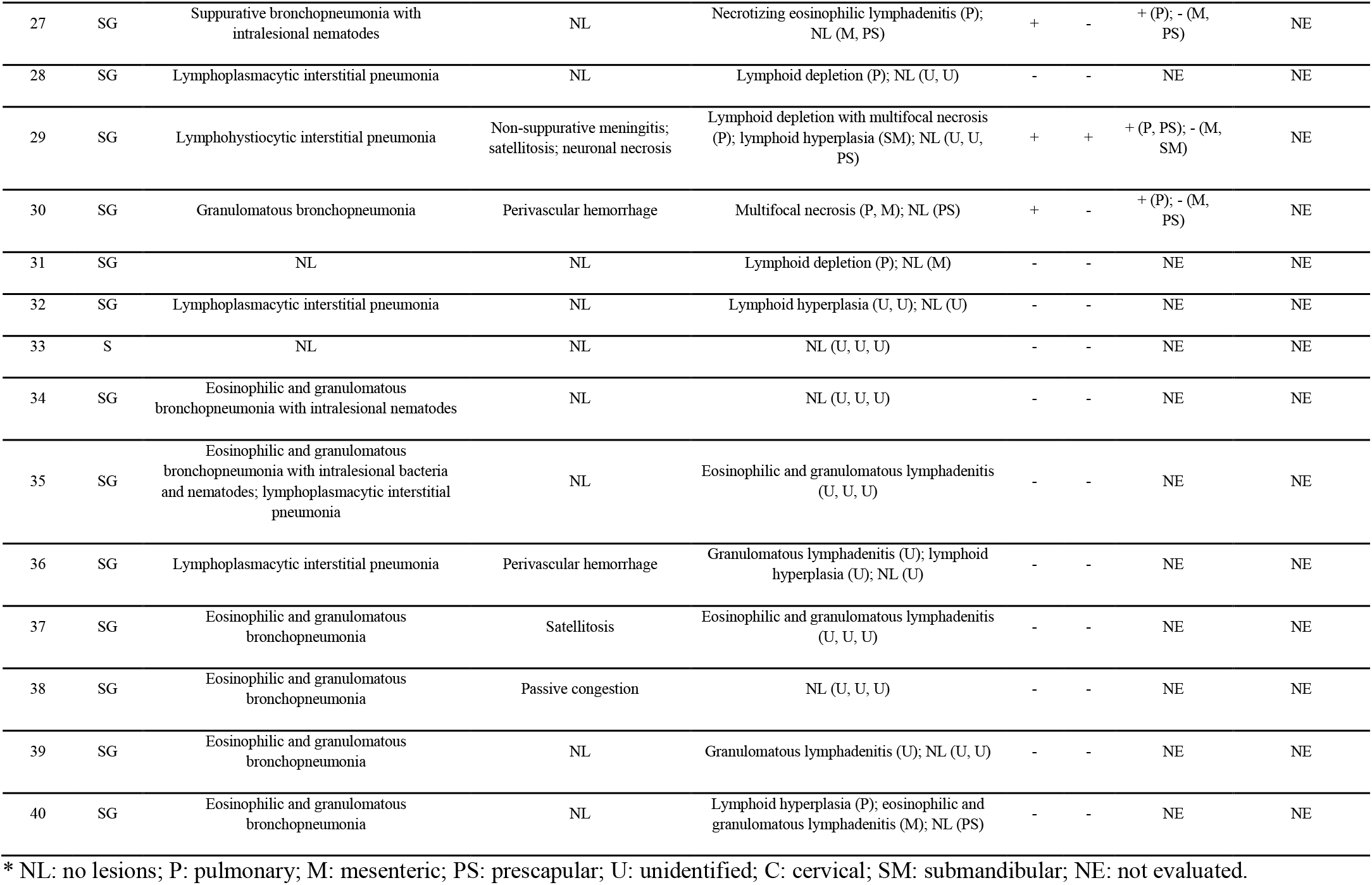

| Animal | Species | Main histopathologic findings |  |  | IHC anti-CeMV |  |  | IHC anti-*Toxoplasma gondii* |
| --- | --- | --- | --- | --- | --- | --- | --- | --- |
|  |  | Lung | CNS | Lymph nodes | Lung | CNS | Lymph nodes | CNS |
| 1 | BA | Pulmonary edema and passive congestion | NL | Diffuse hemorrhage (P, M); NL (PS) | - | - | NE | NE |
| 2 | SG | Lymphoplasmacytic interstitial pneumonia | NL | NL (P, M, PS) | + | - | + (P); - (M, PS) | NE |
| 3 | SG | Lymphoplasmacytic interstitial pneumonia; eosinophilic granulomatous bronchopneumonia | NL | Lymphoid hyperplasia (U); NL (U, U) | + | - | + (U); - (U, U) | NE |
| 4 | SG | Lymphoplasmacytic interstitial pneumonia; eosinophilic granulomatous bronchopneumonia | Neuronal ceroid lipofuscinosis | Lymphoid hyperplasia (U); NL (U, U) | - | - | NE | NE |
| 5 | PB | Granulomatous bronchopneumonia | NL | Focal necrosis (M); NL (P) | - | - | NE | NE |
| 6 | TT | Pulmonary edema | NL | Lymphoid depletion (M); NL (P, PS) | - | - | NE | NE |
| 7 | SG | Granulomatous bronchopneumonia with intralesionais nematodes | NL | Necrotizing granulomatous lymphadenitis with lymphoid depletion (P, M, PS) | + | - | + (P, O); - (M, PS) | NE |
| 8 | SG | Granulomatous bronchopneumonia with intralesionais nematodes | NL | Lymphoid depletion with multifocal necrosis (P, M, PS) | + | - | - (P, M, PS) | NE |
| 9 | TT | Pulmonary edema and hemorrhage | Passive congestion | Lymphoid depletion with mutifocal necrosis (P, M, C, SM); NL (PS) | + | - | + (P, M, C, SM, PS) | NE |
| 10 | PB | Pulmonary edema and passive congestion | NL | Multifocal necrosis (P); lymphoid depletion (PS); NL (SM) | - | - | NE | NE |
| 11 | SG | Lymphoplasmacytic interstitial pneumonia; suppurative bronchopneumonia | Non-suppurative meningoencephalomyelitis with intralesional protozoa cysts | Multifocal necrosis (P, I, M); lymphoid depletion with intracytoplasmic tachyzoites (PS, SM) | - | - | NE | + |
| 12 | SG | Lymphocytic interstitial pneumonia | NL | Necrotizing lymphadenitis with intralesional coccobacillus (P, M) | - | - | NE | NE |
| 13 | SF | Lymphohistiocytic interstitial pneumonia | Non-suppurative encephalitis | Lymphoid depletion (P, M, PS); NL (SM) | - | - | - | NE |
| 14 | SG | Eosinophilic and granulomatous bronchopneumonia with intralesional nematodes | Perivascular hemorrhage | Lymphoid depletion (I); Eosinophilic lymphadenitis (P); NL (SM, PS, M) | - | - | NE | NE |
| 15 | SG | Eosinophilic and granulomatous bronchopneumonia with intralesional nematodes | Perivascular hemorrhage | Lymphoid depletion (P, M, PS, SM) | - | - | NE | NE |
| 16 | SB | Granulomatous bronchopneumonia with intralesional bacteria and nematodes | Passive congestion | Lymphoid depletion (PS, SM, M); Granulomatous lymphadenitis with intralesional parasitic eggs | - | - | NE | NE |
| 17 | SG | Lymphoplasmacytic interstitial pneumonia | NL | NL (P, M, PS, SM) | + | - | + (P, SM); - (M) | NE |
| 18 | SG | Granulomatous bronchopneumonia with intralesional hyphae and bacteria | Non-suppurative encephalitis with intralesional protozoa | Lymphoid depletion (M); NL (P, SM) | - | - | NE | + |
| 19 | SG | Pyogranulomatous pneumonia with intralesional bacteria | NL | Lymphoid hyperplasia (U, U, U) | - | - | NE | NE |
| 20 | SG | Granulomatous bronchopneumonia with intralesional bacteria | Perivascular hemorrhage | Lymphoid depletion (P); lymphoid hyperplasia (C); NL (M, PS) | - | - | NE | NE |
| 21 | PB | Pyogranulomatous pneumonia | Passive congestion | NL (M) | - | - | NE | NE |
| 22 | PB | Passive congestion | NL | NL (U, U) | - | - | NE | NE |
| 23 | SG | Lymphohistiocytic interstitial pneumonia; suppurative bronchopneumonia with intralesional nematodes | Polioencephalomalacia | Multifocal necrosis (P, M, PS) | + | - | + (P); - (M, PS) | NE |
| 24 | SG | Granulomatous bronchopneumonia | NL | Multifocal necrosis (U, U); NL (U) | - | - | NE | NE |
| 25 | SG | Suppurative bronchopneumonia with intralesional nematodes | NL | Lymphoid depletion with multifocal necrosis (U, U) | + | - | + (U, U) | NE |
| 26 | SG | Granulomatous bronchopneumonia with intralesional bacteria | Passive congestion | NL (U, U, U) | - | - | NE | NE |
| 27 | SG | Suppurative bronchopneumonia with intralesional nematodes | NL | Necrotizing eosinophilic lymphadenitis (P); NL (M, PS) | + | - | + (P); - (M, PS) | NE |
| 28 | SG | Lymphoplasmacytic interstitial pneumonia | NL | Lymphoid depletion (P); NL (U, U) | - | - | NE | NE |
| 29 | SG | Lymphohistiocytic interstitial pneumonia | Non-suppurative meningitis; satellitosis; neuronal necrosis | Lymphoid depletion with multifocal necrosis (P); lymphoid hyperplasia (SM); NL (U, U, PS) | + | + | + (P, PS); - (M, SM) | NE |
| 30 | SG | Granulomatous bronchopneumonia | Perivascular hemorrhage | Multifocal necrosis (P, M); NL (PS) | + | - | + (P); - (M, PS) | NE |
| 31 | SG | NL | NL | Lymphoid depletion (P); NL (M) | - | - | NE | NE |
| 32 | SG | Lymphoplasmacytic interstitial pneumonia | NL | Lymphoid hyperplasia (U, U); NL (U) | - | - | NE | NE |
| 33 | S | NL | NL | NL (U, U, U) | - | - | NE | NE |
| 34 | SG | Eosinophilic and granulomatous bronchopneumonia with intralesional nematodes | NL | NL (U, U, U) | - | - | NE | NE |
| 35 | SG | Eosinophilic and granulomatous bronchopneumonia with intralesional bacteria and nematodes; lymphoplasmacytic interstitial pneumonia | NL | Eosinophilic and granulomatous lymphadenitis (U, U, U) | - | - | NE | NE |
| 36 | SG | Lymphoplasmacytic interstitial pneumonia | Perivascular hemorrhage | Granulomatous lymphadenitis (U); lymphoid hyperplasia (U); NL (U) | - | - | NE | NE |
| 37 | SG | Eosinophilic and granulomatous bronchopneumonia | Satellitosis | Eosinophilic and granulomatous lymphadenitis (U, U, U) | - | - | NE | NE |
| 38 | SG | Eosinophilic and granulomatous bronchopneumonia | Passive congestion | NL (U, U, U) | - | - | NE | NE |
| 39 | SG | Eosinophilic and granulomatous bronchopneumonia | NL | Granulomatous lymphadenitis (U); NL (U, U) | - | - | NE | NE |
| 40 | SG | Eosinophilic and granulomatous bronchopneumonia | NL | Lymphoid hyperplasia (P); eosinophilic and granulomatous lymphadenitis (M); NL (PS) | - | - | NE | NE |

